# Dopamine D2 receptors in the bed nucleus of the stria terminalis modulate alcohol-related behaviors

**DOI:** 10.1101/2023.06.13.544820

**Authors:** Dipanwita Pati, Sophia I. Lee, Sara Y. Conley, Tori Sides, Kristen M. Boyt, Avery C. Hunker, Larry S. Zweifel, Thomas L. Kash

## Abstract

Dysregulation of the dopamine (DA) system is a hallmark of substance abuse disorders, including alcohol use disorder (AUD). Of the DA receptor subtypes, the DA D2 receptors (D2Rs) play a key role in the reinforcing effects of alcohol. D2Rs are expressed in numerous brain regions associated with the regulation of appetitive behaviors. One such region is the bed nucleus of the stria terminalis (BNST), which has been linked to the development and maintenance of AUD. Recently, we identified alcohol withdrawal-related neuroadaptations in the periaqueductal gray/dorsal raphe to BNST DA circuit in male mice. However, the role of D2R-expressing BNST neurons in voluntary alcohol consumption is not well characterized. In this study, we used a CRISPR-Cas9-based viral approach, to selectively reduce the expression of D2Rs in BNST VGAT neurons and interrogated the impact of BNST D2Rs in alcohol-related behaviors. In male mice, reduced D2R expression potentiated the stimulatory effects of alcohol and increased voluntary consumption of 20% w/v alcohol in a two-bottle choice intermittent access paradigm. This effect was not specific to alcohol, as D2R deletion also increased sucrose intake in male mice. Interestingly, cell-specific deletion of BNST D2Rs in female mice did not alter alcohol-related behaviors but lowered the threshold for mechanical pain sensitivity. Collectively, our findings suggest a role for postsynaptic BNST D2Rs in the modulation of sex-specific behavioral responses to alcohol and sucrose.

## Introduction

Alcohol use disorder (AUD) is the most prevalent substance use disorder worldwide. According to a recent study, the rate of alcohol-related deaths increased to ∼25% during the first year of the COVID-19 pandemic (White et al., 2022). AUD is characterized by repeated cycles of drinking and withdrawal accompanied by neuroadaptations in brain regions involved in motivated behaviors (Koob and Volkow, 2010). Decades of research have implicated several neurotransmitter systems in the pathophysiology of AUD including the mesocorticolimbic dopamine (DA) system (Volkow et al., 2009; Koob and Volkow, 2016). Alcohol increases the firing rate of ventral tegmental area (VTA) DA neurons (Brodie et al., 1990) along with an increase in dopamine release in several brain regions, such as the nucleus accumbens (Di Chiara and Imperato, 1988; Boileau et al., 2003) and striatum (Melendez et al., 2003). However, repeated cycles of withdrawal from chronic alcohol result in reduced DA release, and this hypodopaminergic state is hypothesized to drive craving (Rossetti et al., 1992; Diana et al., 1993; Weiss et al., 1996).

DA acts on two main subfamilies of receptors, D1 and D2. These subfamilies can be further divided into subtypes, with the D1-like subfamily including dopamine D1 and D5 receptors and the D2-like subfamily including dopamine D2, D3, and D4 receptors (Beaulieu and Gainetdinov, 2011). Several lines of evidence from both human and animal studies suggest the importance of D2 receptors in AUD. Reduced levels of D2Rs in limbic areas have been observed in both AUD patients (Hietala et al., 1994; Volkow et al., 1996, 2002, 2013; Tupala et al., 2001) and in alcohol-preferring rodents (STEFANINI et al., 1992; McBride et al., 1993; Feltmann et al., 2018). However, findings from preclinical studies in animal models have been inconsistent. For example, while pharmacological blockade of D2-like receptors in NAc reduced ethanol seeking and drinking (Kaczmarek and Kiefer, 2000; Czachowski et al., 2001), overexpression of striatal D2Rs also reduced ethanol self-administration and preference (Thanos et al., 2001).

One possible reason for these confounding data is that D2Rs have differential effects on alcohol consumption in a cell-type and region-specific manner. However, to date, most studies have focused on ventral tegmental area (VTA) DA neurons and their striatal projections. The bed nucleus of the stria terminalis (BNST) is an important hub interconnecting stress and reward circuits (Erb et al., 2001; Aston-Jones and Harris, 2004; Vranjkovic et al., 2017) and receives dopaminergic inputs from several midbrain regions, including VTA, and ventrolateral periaqueductal gray/dorsal raphe (vlPAG/DR) (Freedman and Cassell, 1994; Hasue and Shammah-Lagnado, 2002; Li et al., 2016). Additionally, the vast majority of neurons in the dBNST are GABAergic in phenotype (Sun and Cassell, 1993) and send projections to areas implicated in AUD, including the VTA (Dong et al., 2001; Georges and Aston-Jones, 2002; Dong and Swanson, 2006). Similar to the nucleus accumbens, alcohol dose-dependently increased extracellular BNST DA concentrations (Carboni et al., 2000). Further, the existence of D2Rs in BNST has been characterized using a combination of slice physiology (Krawczyk et al., 2011b, 2011a; Li et al., 2016; Yu et al., 2021b), in situ hybridization (De Bundel et al., 2016; Melchior et al., 2021; Yu et al., 2021b) and receptor autoradiography (Bouthenet et al., 1987) approaches. Prior studies have implicated BNST D2Rs in the modulation of fear generalization (De Bundel et al., 2016) and impulsivity (Kim et al., 2018). However, the role of BNST D2Rs in alcohol consumption remains unclear. Here, we used a CRISPR-Cas9-based viral approach to selectively knockout D2Rs from BNST GABA neurons and explored the role of BNST D2Rs in alcohol-related behaviors in both male and female mice.

## Materials and Methods

### Mice

Dopamine transporter expressing (DAT-cre) mice (Zhuang et al., 2005) were bred and housed at the University of Washington, Seattle. Vesicular GABA transporter expressing (VGAT-ires-Cre) mice were generated as previously described (Krashes et al., 2014), were bred in-house, and maintained on a C57BL6/J background. All behavioral experiments were performed on adult male (N=33) and female mice (N=41), aged 8-16 weeks at the start of the procedures, in accordance with the NIH guidelines for animal research and with the approval of the Institutional Animal Care and Use Committee at the University of North Carolina at Chapel Hill. For all behavioral experiments, except the drinking studies, mice were maintained on a standard 12-hour cycle (lights on at 07:00) in a temperature-controlled colony with *ad libitum* access to food and water. For the drinking experiments, mice were transferred to a reverse light cycle (light off at 07:00) and allowed to acclimate for a week before the start of the experiments.

### Design of *Drd2* CRISPR construct

A single guide RNA (sgRNA) targeting *Drd2* was designed as described before (Star Protocols; Hunker et al., 2020). Primers for sgDrd2 were: forward CACCGAGAACTGGAGCCGGCCCTTCA and reverse AAACTGAAGGGCCGGCTCCAGTTCTC. AAV1-FLEX-SaCas9-U6-sgRosa26 was previously published (Hunker et al., 2020). AAVs for CRISPR constructs were purified as described (Hunker et al., 2020). For sequencing validation, AAV1-FLEX-SaCas9-sgDrd2 was injected into the VTA of DAT-Cre mice along with AAV1-FLEX-EGFP-KASH. Four weeks following injection the VTA was micro-dissected, nuclei were isolated and fluorescently sorted as described in Hunker et al., 2020. The sequence flanking the sgRNA target was amplified by PCR with the following primers: initial forward: TCTCAGCTCTGCTAGCTC and initial reverse: GCATAACCAGTGTGGCCA; final forward: ACTCCTGCTCACTCCTG and final reverse: CTTCTCTCTGGATACAGC. The final amplicon was deep sequencing using Amplicon-EZ (Genewiz).

### Surgical procedure

Adult mice were anesthetized with isoflurane (2%–3%) in oxygen (0.8-1 l/min) and then secured on a stereotaxic frame (Model 1900, Kopf Instruments, CA) for intracranial viral infusions. To minimize postoperative discomfort, meloxicam (5mg/kg) was administered at the time of the surgery and for two additional days. The BNST (from bregma in mm: AP +0.14 mm, ML ±0.90 mm, DV -4.25 mm) was targeted using standard coordinates from Paxinos & Franklin. Microinjections were performed with a 1 μl Neuros Hamilton syringe (Hamilton, Reno, NV) controlled by a micro-infusion pump (WPI, FL) at a rate of 100 nl/min. For behavioral experiments, AAV1-CMV-FLEX-saCas9-U6-sgDrd2 (plasmid gift from Dr. Larry Zweifel, packaged at UNC vector core; 1.3×10^12^ vg/mL) was mixed with AAV1-CAG-FLEX-EGFP-WPRE (Addgene; Cat # 51502; ≥ 1×10^13^ vg/mL) in a 4:1 ratio. 300 nl of this mixture was infused bilaterally in the BNST of VGAT-cre mice. The control mice received a mixture of AAV1-CMV-FLEX-saCas9-U6-sgRosa26 (plasmid gift from Dr. Larry Zweifel, packaged at UNC vector core; 3.8×10^12^ vg/mL) and the cre-dependent GFP virus. For the validation of the CRISPR-Cas9-mediated deletion of D2 receptors, a separate cohort of mice was infused with the viral mixture in a 3:1 ratio. After infusion, injectors were left in place for 5 min to allow for viral diffusion. All surgeries were conducted using aseptic techniques in a sterile environment. Mice were closely monitored and allowed to recover before starting experiments.

### Behavioral assays

∼3-4 weeks post-surgery, age-matched mice were randomly assigned to experimental groups. One week before the first behavioral test, mice were singly housed and maintained on an Isopro RMH 3000 (LabDiet, St. Louis, MO, USA) diet, which has been shown to produce high levels of alcohol consumption (Marshall et al., 2015). On the day of the tests, animals were transported to the behavioral room and allowed to acclimate to the room for ∼45-60 minutes. The experimental timeline is shown in Fig. 2. All mice went through the same sequence of behavioral tests. All behavioral testing except the drinking study, was carried out during the light cycle, and there was at least 48 h between test sessions. Male and female mice were tested on different days except for the ethanol-induced locomotor assay. The experimenter was blind to the genotype of the animals throughout behavioral experiments and data analysis.

### Open Field

The open field arena consisted of an apparatus made of white Plexiglas (50 × 50 × 25 cm). Mice were placed into the corner of the arena and allowed to freely explore for 10 min. The center of the open field was defined as the central 25% of the arena. Light levels were ∼200 lux. Sessions were video recorded and analyzed using the EthoVision software (Noldus Information Technologies, Leesburg, VA, USA).

### Mechanical sensitivity

The Von Frey test was used to measure mechanical nociception, as described before (Yu et al., 2021b). Briefly, mice were confined to Plexiglas boxes on an elevated metal wire surface (90 × 20 × 30 cm) and habituated to the behavioral apparatus for a minimum of 60 minutes. Nylon monofilaments of forces ranging from 0.008 to 2 g (g) were applied to the hind paws using the simplified up-down method (SUDO) (Bonin et al., 2014). Starting with a mid-range force (0.16 g), the filament was applied to the mid-plantar surface of the hind paw for ten trials, then repeated with ascending or descending forces depending on the number of paw withdrawals. Withdrawal thresholds were defined as the minimum force (g) filament that elicits a withdrawal reflex for ≥ 50% of the trials.

### Thermal sensitivity

Thermal sensitivity was measured using the Hargreaves test (Hargreaves et al., 1988) as described before (Yu et al., 2021b). Mice were placed in Plexiglas boxes on an elevated glass surface and habituated to the behavioral apparatus for at least 60 minutes. The beam intensity was set on the IITC Plantar Analgesia Meter (IITC Life Science, Woodland Hills, CA). The mid-plantar surface of each hind paw was then exposed to a series of heat trials with 10-minute inter-trial intervals. Four trials were conducted with radiant heat exposures sequentially alternating between left and right paws. Paw withdrawal latency was recorded. A cutoff time of 25 s was set to prevent excessive tissue damage.

### Acoustic startle

The acoustic startle test was performed with the SR-LAB Startle Response System (SD Instruments). The system consisted of a sound-attenuating isolation cabinet with dimensions 38 cm × 36 cm × 46 cm containing a small plastic cylindrical enclosure with dimensions 4 cm (diameter) × 13 cm (length). Mice were placed in the plastic enclosure and allowed to acclimate for 5 min. After acclimation, mice were presented with the first of the startle stimulus trials. Ten stimuli of each intensity level (0, 90, 105, and 120 dB) were presented in a pseudo-random order (the constraint being that each intensity occurs within each block of four trials) with an intertrial interval of 30-50s. Stimuli were delivered for 40 ms in each trial and the startle response was measured for 200 ms following stimulus delivery. Startle amplitude was defined as the largest peak 100 ms after the onset of the startle stimulus. For each animal, startle amplitudes for each stimulus level were averaged across the ten trials.

### Ethanol-induced locomotion

Ethanol-induced locomotor activity was measured using a SuperFlex open field system (Omnitech Electronics, Accuscan, Columbus, OH), and a proprietary Fusion v6.5 software was used to track beam breaks to determine the total distance traveled during 30 min testing sessions (Fig. 5A). The open field arena measured 60 cm × 60 cm × 40 cm and the light levels were∼25-30 l×. To reduce injection stress and novelty effects, mice were injected intraperitoneally (IP) with saline and were habituated to the locomotor boxes on days 1 and 2. On days 3 and 4, mice were placed in the locomotor boxes right after IP injection of saline or ethanol (2g/kg) in a counterbalanced, crossover design. Ethanol (95%) was prepared 20% v/v in a solution of 0.9% saline and administered intraperitoneally (IP) at a dose of 2 g/kg in a 12.5 mL/kg volume. Distance traveled (cm) was measured in 5 min bins for 30 min during the two testing sessions.

### 2-Bottle choice intermittent access to ethanol (2-BC IA)

After the completion of the last behavioral experiment, mice were transferred to the reverse light cycle and were habituated to two 50 ml water bottles. One week later, both male and female mice underwent 6 weeks of 2-BC IA (Fig. 6A), as described before(Bloodgood et al., 2020). Briefly, ∼3 h into the dark cycle, mice were given free access to both water and 20% w/v ethanol bottles in their home cage for 24 h. Drinking sessions occurred three times per week (Mon, Wed, Fri). Bottles were weighed at the start and end of each session. At all other times, mice were given access to water alone. Ethanol and water bottle positions were alternated each session to account for any inherent side bias. This weekly access schedule was repeated for 6 weeks for a total of 18 sessions. To account for the fluid lost due to passive drip, ethanol, and water bottles were placed in an empty cage and weighed during each drinking session. Drip values were then calculated for each solution and total consumption was normalized to these values. The water mice had access to water alone. All mice were weighed every week and ethanol intake was calculated as g/kg/24h. 20% w/v ethanol was prepared fresh each week from 190-proof ethanol mixed with tap water.

### Voluntary sucrose consumption

In one cohort of mice, 5 days after the final IA session, both water and ethanol mice were given free access to both water and 0.5% w/v sucrose bottles in their home cage for 2 h a day, 3 h into the dark cycle, for 5 days. Sucrose and water bottle positions were alternated each session to account for any inherent side bias. The drinking values for the last three sessions were averaged and compared.

### Histology

All mice used for behavioral experiments were anesthetized with tribromoethanol (Avertin, 1 ml, IP) and transcardially perfused with ice-cold 0.01 M phosphate-buffered saline (PBS) followed by 4% paraformaldehyde (PFA) in PBS. Brains were extracted and stored in 4% PFA for 24 hours followed by stored in 30% sucrose/PBS at 4 °C until the brains sank. 45 μm coronal sections were collected using a Leica VT1000S vibratome (Leica Biosystems; Deer Park, IL, USA) and then stored in PBS with 0.02% sodium azide (Sigma Aldrich). Sections were mounted on slides, allowed to dry, coverslipped with VectaShield Hardset Mounting Medium with DAPI (Vector Labs, Newark, CA, USA), and stored in the dark at 4 °C. Keyence BZ-X800 All-in-one Fluorescence microscope (Keyence; Itasca, IL, USA) was used to image GFP expression to verify viral injection sites (Fig. 1F-G).

**Figure 1.**
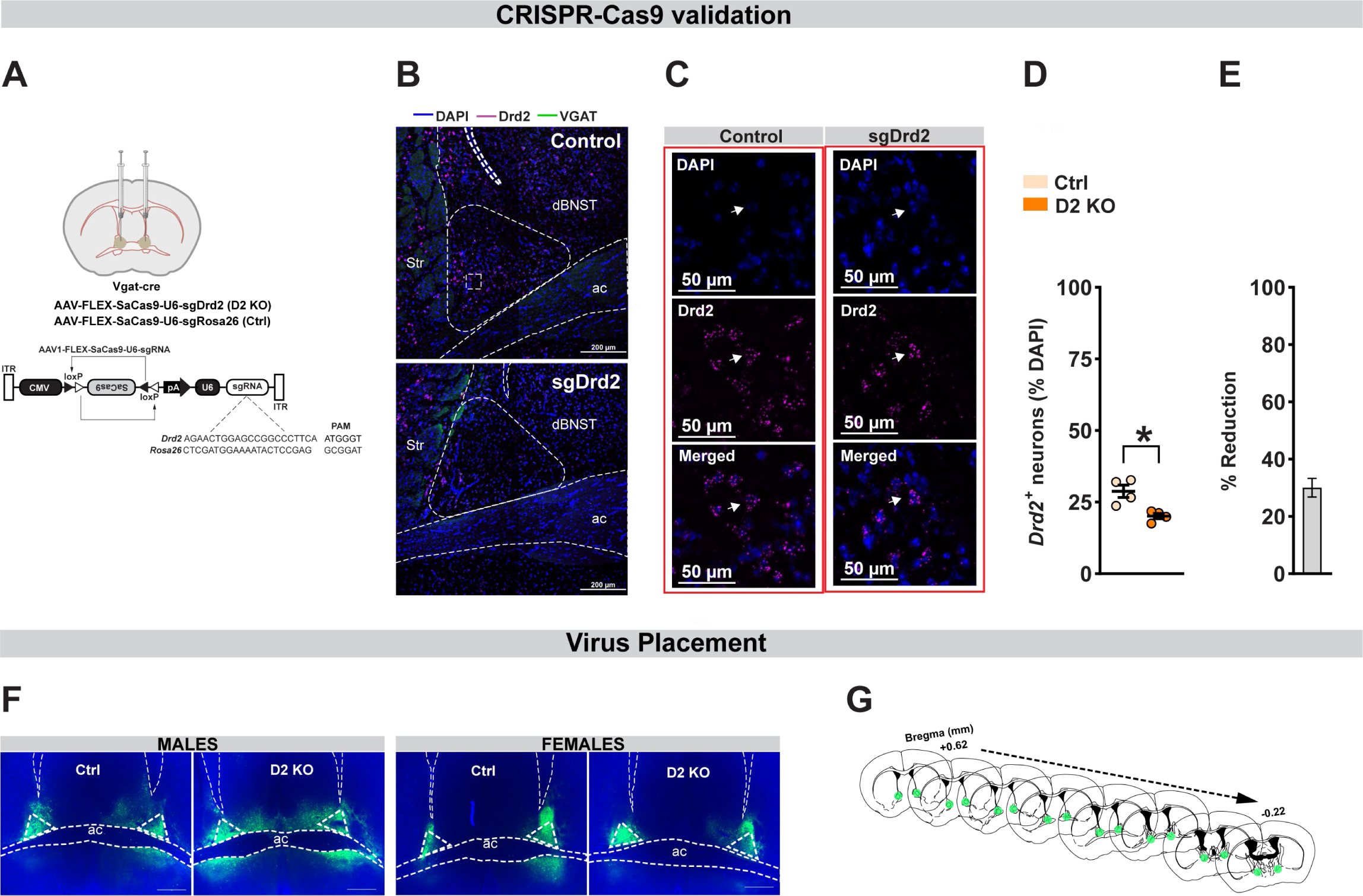
Validation of CRISPR-Cas9 deletion of BNST D2Rs. **(A)** Top: Surgical schematic of bilateral virus infusion in the BNST of VGAT-cre male and female mice. Bottom: Detailed schematic of *Drd2* (D2 KO) and Rosa26 (control) CRISPR construct provided by the Zweifel lab. **(B)** Representative images of in situ hybridization for D2 (cyan) and VGAT (green) mRNA from control (top) and D2 KO (bottom) in the mouse BNST. Scale bar=200 μm. **(C)** Representative images at higher magnification showing *Drd2* (cyan) labeling inside neurons labeled with DAPI (blue) from control (left) and D2 KO (right) mice. Scale bar=50 μm. **(D)** Compared to controls, D2 KO mice showed a significant reduction in *Drd2* mRNA expression in the dBNST. **(E)** CRISPR knockout resulted in ∼30% reduction in *Drd2*+ mRNA expression. **(F)** Representative viral infections in BNST in males (left) and females (right), as indexed by GFP expression. Scale bar=500 μm. **(G)** Representative map of viral spread from +0.62 mm to -0.22 mm (from bregma). ac: anterior commissure; str: striatum. Data expressed as Mean ± SEM. *p<0.05.

### Fluorescent in situ hybridization (FISH)

To validate *Drd2* deletion, mice were anesthetized with isoflurane, rapidly decapitated, and brains were collected and immediately frozen on dry ice and stored in a −80°C freezer. 12 µm thick coronal sections of BNST were obtained using a Leica CM3050 S cryostat (Leica Microsystems, Wetzlar, Germany), and directly mounted onto Superfrost Plus slides (Fisher Scientific, Hampton, NH), and stored at -80°C. RNAscope was performed according to the manufacturer’s instructions (Advanced Cell Diagnostics, Newark, CA). To fluorescently label *Drd2* and VGAT mRNA in the BNST, slices were preprocessed with 4% PFA and protease reagent, incubated with target probes for mouse *Drd2* and VGAT, then fluorescently labeled with probes targeting the corresponding channels (VGAT 488; Drd2 647; Advanced Cell Diagnostics, Newark, CA). The processed slides were then coverslipped using ProLong Gold Antifade Mountant with DAPI.

### Confocal microscopy

Images were captured using a Zeiss 800 upright confocal microscope (Hooker Imaging Core, UNC-Chapel Hill). For quantification of D2R knockout, tiled and serial z-stack images were obtained through a 20x objective (2 μm optical slice thickness), and converted to maximum projection intensity images using Zen blue software (2 slices/animal). Analysis was performed using subcellular particle detection in QuPath (Bankhead et al., 2017). Cells were considered positive if they contained ≥ 5 puncta of *Drd2* mRNA.

### Data and statistical analysis

All statistical analysis was performed using GraphPad Prism v.9 (La Jolla, CA, USA). Group comparisons were made using either a standard unpaired Student’s t-test or two/three-way ANOVA depending on the number of independent and within-subjects variables in a dataset. Following significant interactions or main effects, planned post hoc analyses were performed and corrected for multiplicity. Male and female data were analyzed disaggregated as well as with a factorial design with sex as a covariate. If data violated assumptions of normality a Mann-Whitney U test was used for analysis. Data are expressed as mean ± SEM unless otherwise stated. P-values ≤ 0.05 were considered significant. A total of 4 mice were excluded from the analysis due to no AAV expression within the BNST.

## Results

### CRISPR-Cas9-mediated deletion of dopamine D2Rs

To define the contribution of BNST D2 receptors in alcohol-related behaviors, we used a CRISPR-Cas9 system to conditionally knock out D2Rs from BNST^VGAT^ neurons (D2 KO) in male and female mice. We packaged a plasmid for cre-dependent expression of *Staphylococcus aureus* Cas9 (SaCas9) plus a single guide RNA (sgRNA) targeted to the gene encoding *Drd2* (sgDrd2) in an adeno-associated virus (AAV) vector (Fig. 1A) (Hunker et al., 2020). Mutagenesis was confirmed by targeted deep sequencing of the genomic region flanking the sgRNA (Supplementary to Fig.1). In a separate cohort, we injected sgDrd2 construct or a control virus (sgRosa26) bilaterally into the BNST of VGAT-cre mice (N=2 mice/sex/group). An additional cre-dependent virus that drove the expression of GFP was co-administered to visualize the injection site. After waiting for at least 4 weeks to ensure sufficient viral expression we confirmed the mutagenesis of *Drd2* by performing fluorescent in situ hybridization (Fig. 1B-C).

Quantification of *Drd2* mRNA expression in dBNST revealed a significant reduction in D2 KO mice compared to control animals [Fig. 1D; t_6_ = 3.64; p = 0.011]. When normalized to average *Drd2*^+^ neurons in the control group, CRISPR knockout resulted in ∼30% reduction in *Drd2*+ mRNA expression [Fig. 1E; t_3_ = 9.19; p = 0.003; one sample t test]. In mice used for behavioral testing, we confirmed viral infection in BNST using fluorescence microscopy to visualize GFP expression (Fig. 1F-G).

### Effect of BNST D2R deletion on open field behavior and pain sensitivity

After waiting for 4 weeks to ensure sufficient expression of the virus, all mice went through a battery of behavioral assays to assess the impact of BNST D2R loss of function on anxiety-like, pain sensitivity, and alcohol-related behaviors (Fig. 2). To assess the effect of BNST^VGAT^ D2R deletion on anxiety-like behavior, mice were placed in an open field arena and the total distance traveled and the time spent in the center zone was calculated. Selective knockout of D2Rs (N=15-16 male and 19-20 female mice per group) did not affect total distance traveled (Fig. 3A). A two-way ANOVA revealed a significant main effect of sex [F_(1,66)_=8.723; p_sex_ =0.004] but no main effect of virus [F_(1,66)_=0.627; p_virus_=0.431] or sex x virus interaction [F_(1,66)_=0.0003; p_interaction_=0.987]. Post-hoc comparisons showed a trend for increased locomotion in females [t_66_ = 2.13, p_corrected_ = 0.072 for control mice and t_66_ = 2.05, p_corrected_ = 0.087; Sidak’s test]. Deletion of D2Rs did not alter the time spent in the center zone in both sexes (Fig. 3B; [F_(1,66)_=0.048; p_virus_ =0.827]; [F_(1,66)_=0.676; p_sex_ =0.414]; [F_(1,66)_=0.322; p_interaction_ =0.572]).

**Figure 2.**
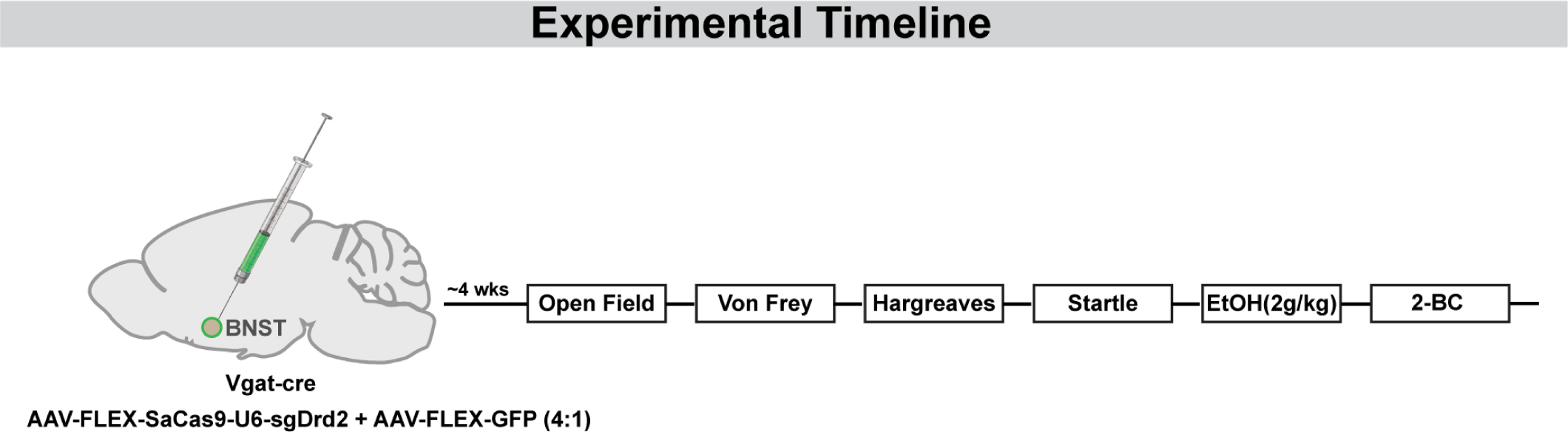
Experimental Timeline. Schematics of the experimental timeline showing the sequence followed for behavioral assays in all cohorts of mice.

**Figure 3.**
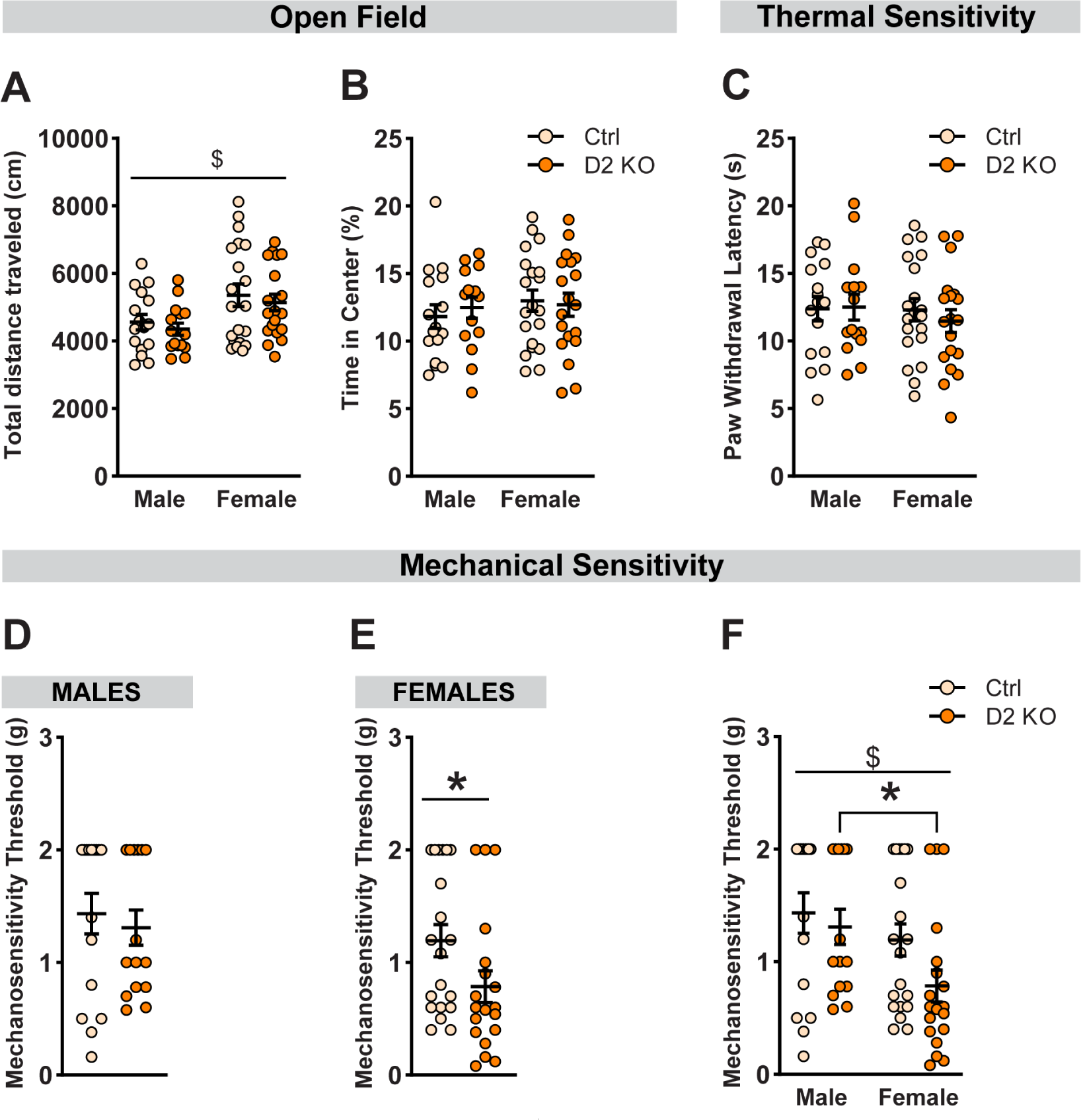
Impact of BNST D2R deletion on anxiety-like behavior and pain sensitivity. Anxiety-like behavior was assessed in an open field assay. No differences were observed either in the total distance traveled (cm) **(A)** or percentage time spent in the center of the open field arena **(B)** following BNST D2R knockout. Hargreaves test was used to assess thermal sensitivity and average paw withdrawal latencies were compared **(C)**. In both sexes, deletion of D2-Rs had no impact on withdrawal latency. Von Frey test was used to assess the contribution of BNST D2Rs in mechanical sensitivity **(D-F)**. Paw withdrawal threshold was defined as the minimum force (g) filament that elicits a withdrawal reflex for ≥ 50% of the trials. **(D)** No effect of D2 knockout on withdrawal threshold in male mice. **(E)** D2 KO female mice had a significantly lower withdrawal threshold compared to female controls. **(F)** There was a main effect of sex on mechanical pain threshold and a non-significant trend for a main effect of deletion. Post-hoc analysis revealed a significantly lower threshold in D2 KO females compared to D2 KO males. Data expressed as Mean ± SEM.*p<0.05; $ main effect of sex.

Next, we asked whether a cell-type-specific reduction in D2Rs could modulate pain sensitivity. The Hargreaves test was used to measure thermal sensitivity. We found no significant effect of knockout on paw withdrawal latency in both sexes (Fig. 3C; [F_(1,66)_=0.165; p_virus_ =0.686]; [F_(1,66)_=0.39; p_sex_ =0.535]; [F_(1,66)_=0.286; p_interaction_ =0.594]). To assess the impact of D2R deletion on mechanical sensitivity, we used the Von Frey assay. Interestingly, when analyzed separately, D2 KO female mice showed a significant reduction in withdrawal threshold (Fig. 3E; [U = 114.5; p = 0.032]). This effect was absent in male mice (Fig. 3D; p=0.608). A two-way ANOVA with sex and virus as between-subject factors (Fig. 3F) showed a significant main effect of sex [F_(1,66)_=6.038; p_sex_ =0.017] and a non-significant trend for knockout effect [F_(1,66)_=2.94; p_virus_=0.091], but no sex x virus interaction [F_(1,66)_=0.837; p_interaction_=0.364]. Importantly, post-hoc analyses showed a significant difference in mechanical sensitivity in D2 KO females compared to D2 KO males [U = 66.5, p_corrected_ = 0.013; Holm-Sidak test].

### Reduction in BNST D2Rs does not alter startle response

To characterize the role of BNST D2Rs in regulating arousal-like behavior, mice were tested for acoustic startle responses (Fig. 4). Male (N=15-16/group) and female mice (N=19-20/group) were presented with four different levels of sound intensities in a randomized manner and the startle amplitudes were recorded. Sound intensities ranging from 0-120 dB resulted in increased startle responses in both males (Fig. 4A; [F_(1.7,48.17)_=44.26; p_dB_<0.001]) and females (Fig. 4B; [F_(1.71,63.27)_=53.32; p_dB_<0.001]) as revealed by repeated-measures two-way ANOVA with Geisser-Greenhouse correction. However, there was no main effect of D2 knockout ([F_(1,29)_=0.104; p_virus_=0.75 for males]; [F_(1,37)_=0.186; p_virus_=0.67 for females]) or dB x virus interaction ([F_(3,87)_=0.294; p_interaction_=0.83 for males]; [F_(3,111)_=1.91; p_interaction_=0.133 for females]) in both sexes. Comparing the startle amplitude at 120 dB (Fig. 4C) revealed a main effect of sex [F_(1,66)_=12.5; p_sex_=0.0007], but no effect of virus [F_(1,66)_=0.047; p_virus_=0.83] or sex x virus interaction [F_(1,66)_=1.08; p_interaction_=0.303]. Post-hoc analyses with Holm-Sidak correction indicated an overall reduced startle response in females compared to males ([U= 86, p_corrected_ = 0.036; for controls]; [U= 77, p_corrected_ = 0.036; for D2 KOs]).

**Figure 4.**
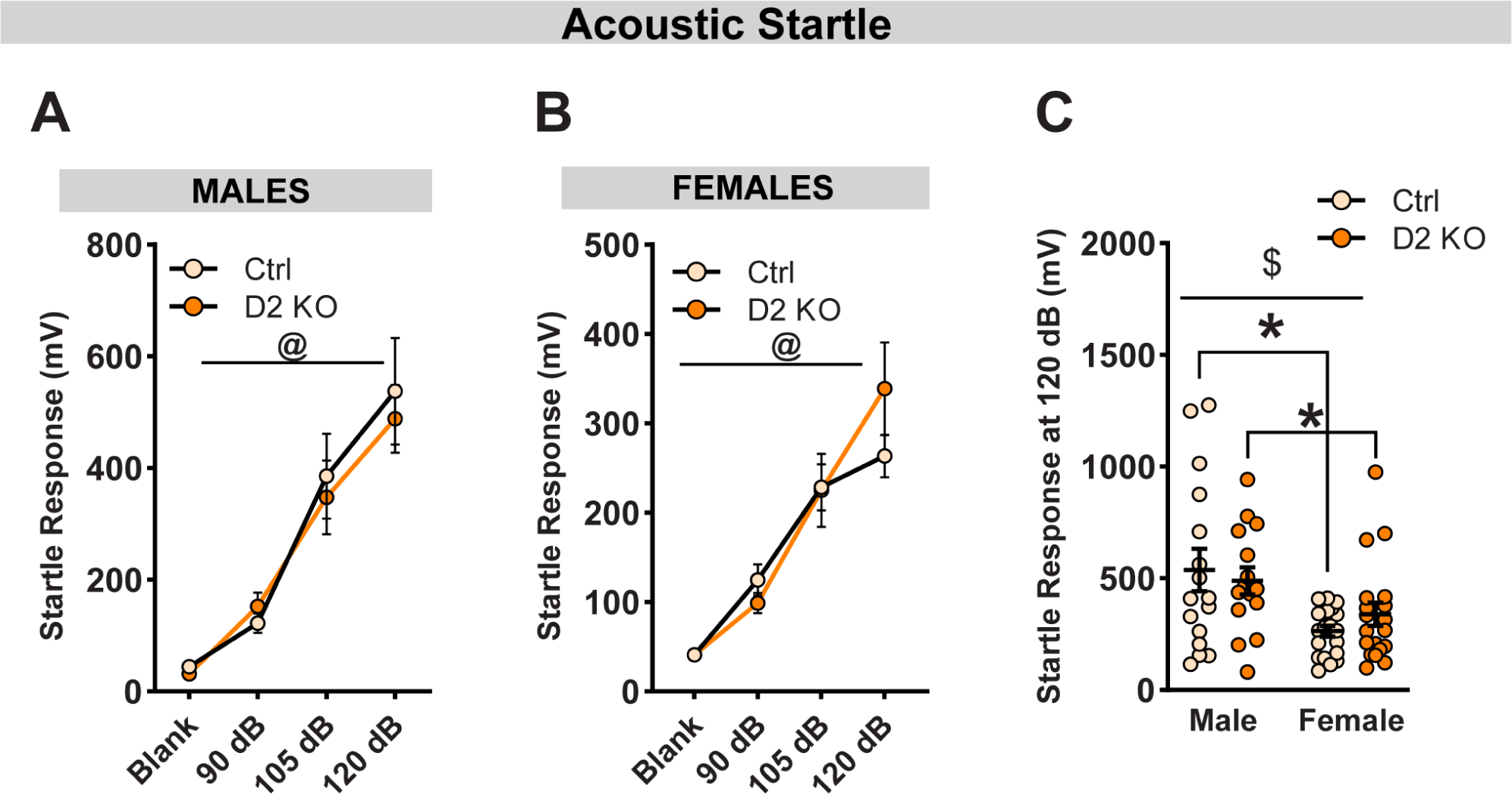
BNST^VGAT^ D2R knockout does not affect acoustic startle response. Average startle amplitudes (10 trials/dB) were plotted in response to increasing sound intensities. No effect of reduced D2R expression on startle amplitude in males **(A)** and females **(B)**. **(C)** Comparison of startle response at 120 dB between male and female mice. Overall, female mice displayed lower amplitude compared to male mice. No differences were observed between D2 KO and control mice. Data expressed as Mean ± SEM.*p<0.05; @ main effect of decibel; $ main effect of sex.

### Selective loss of D2Rs enhances stimulatory effects of ethanol in a sex-specific manner

Global knockout of D2 receptors in mice enhances ethanol-induced locomotion (Palmer et al., 2003) in a familiar environment. Therefore, we wanted to evaluate whether deletion of D2Rs from BNST VGAT neurons altered the stimulatory effects of acute ethanol (Fig. 5). After habituation to the locomotor boxes, male (N=14-15/group) and female (N=19/group) mice were challenged with saline and 2g/kg of ethanol in a counterbalanced, within-subjects design, and locomotion was quantified using infrared beam breaks (Fig. 5A). Consistent with the literature, ethanol produced transient, stimulatory effects that peaked within the first 5 minutes (Fig. 5B-C) (Liljequist et al., 1981; Cohen et al., 1997). When both sexes were analyzed separately, a three-way RM-ANOVA revealed a main effect of time ([F_(5,135)_=60.54; p_time_<0.0001 for males]; [F_(5,180)_=220.5; p_time_<0.0001 for females]), ethanol ([F_(1,27)_=26.07; p_etoh_<0.001 for males]; [F_(1,36)_=15.31; p_etoh_=0.004 for females]) and a time x ethanol interaction ([F_(5,135)_=26.91; p_interaction_<0.0001 for males]; [F_(5,180)_=108.4; p_interaction_<0.0001 for females]). Interestingly, only male mice had a significant effect of virus ([F_(1,27)_=5.59; p_virus_=0.026 for males]; [F_(1,36)_=0.004; p_virus_=0.947 for females]). A two-way RM-ANOVA in male mice comparing the distance traveled during the first 5 minutes resulted in heightened ethanol-induced stimulation in D2 KO mice (Fig. 5D). Specifically, there was a main effect of both virus [F_(1,27)_=4.83; p_virus_=0.037] and ethanol [F_(1,27)_=68.54; p_etoh_<0.001], but no interaction effects [F_(1,27)_=2.08; p_interaction_=0.16]. Post-hoc analyses with Holm-Sidak correction indicated a significant increase in locomotion following ethanol injection when compared to saline ([t_27_=4.92, p_corrected_<0.0001; for controls]; [t_27_=6.76, p_corrected_<0.0001; for D2 KOs]) with the magnitude of response being significantly larger in D2 KO male mice [t_54_=2.63, p_corrected_ = 0.022]. In female mice (Fig. 5E), there was a main effect of ethanol [F_(1,36)_=221.6; p_etoh_<0.001], but no effect of virus [F_(1,36)_=1.01; p_virus_=0.322] or a virus x etoh interaction [F_(1,36)_=1.66; p_interaction_=0.205]. Similar to male mice, post-hoc analyses indicated a significant increase in locomotion following ethanol injection when compared to saline ([t_36_=11.44, p_corrected_<0.0001; for controls]; [t_36_=9.62, p_corrected_<0.0001; for D2 KOs]; Holm-Sidak correction). When compared with sex as a covariate (Fig. 5F), a 3-way RM-ANOVA revealed a main effect of ethanol [F_(1,63)_=257.1; p_etoh_<0.0001], sex [F_(1,63)_=20.53; p_sex_<0.0001] and significant interactions between ethanol x sex [F_(1,63)_=13.24; p_interaction_=0.0006] and sex x virus [F_(1,63)_=5.01; p_interaction_=0.029]. Additionally, there was a statistical trend toward a 3-way interaction of sex, virus, and ethanol [F_(1,63)_=3.74; p_interaction_=0.058]. Consistent with known sex differences (Crabbe et al., 1987), on average, control female mice showed higher ethanol-induced stimulation when compared to male mice ([t_126_=6.24, p_corrected_<0.0001]; Holm-Sidak’s correction). Compared to saline, ethanol injection resulted in increased locomotion in both sexes across groups (all p<0.0001).

**Figure 5.**
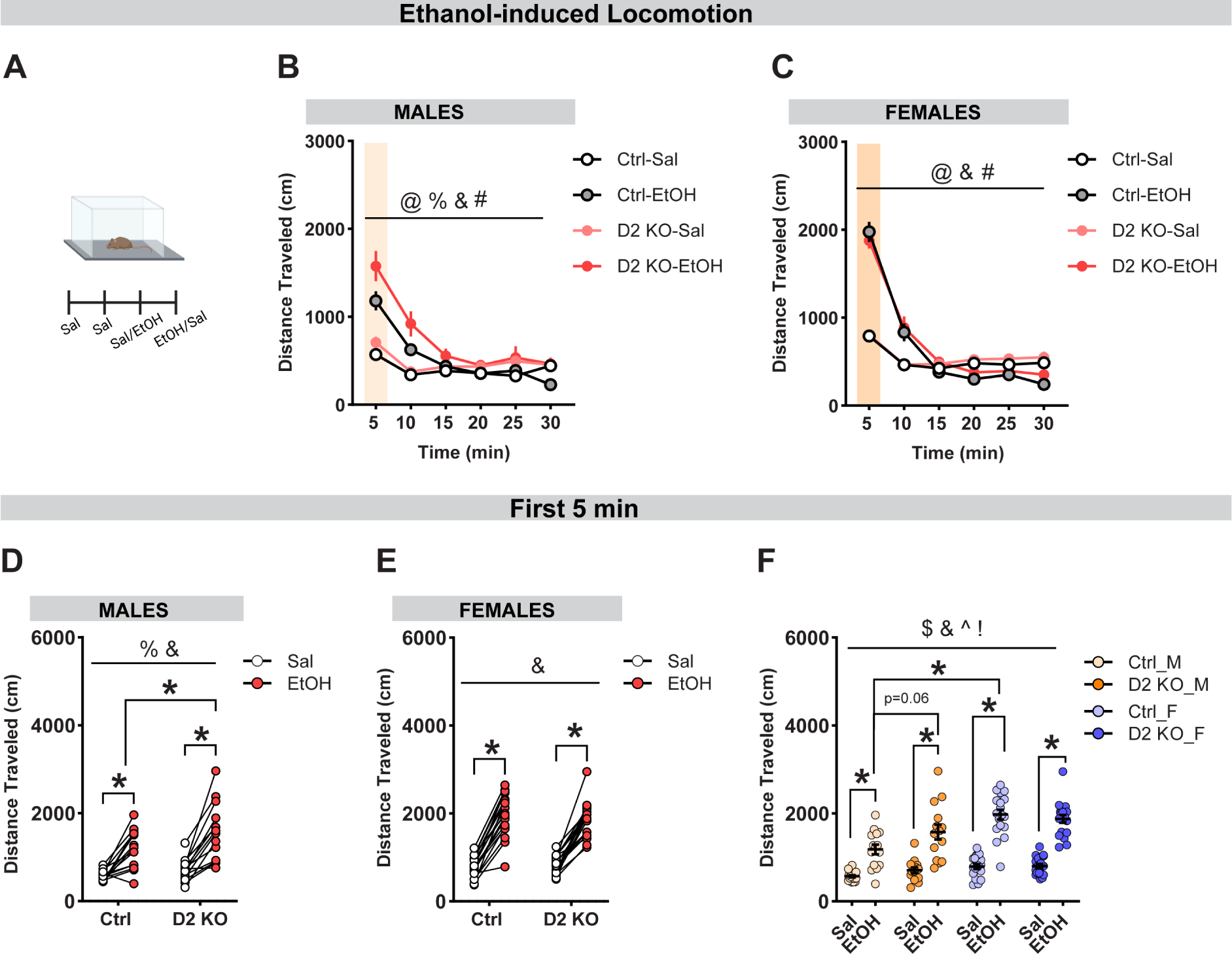
Selective deletion of BNST^VGAT^ D2Rs promotes ethanol-induced locomotion in male mice. **(A)** Schematic diagram representing ethanol-induced locomotor assay. Mice were habituated to the locomotor boxes for two days followed by IP injection of saline and 2g/kg ethanol in a counter-balanced, crossover design. Total distance traveled (cm) was plotted in 5-minute bins for 30 minutes. 2g/kg ethanol injection resulted in a transient increase in locomotion in both sexes **(B-C)**. Deletion of D2Rs altered the total distance traveled in a sex-dependent manner. **(D-F)** Average distance traveled in the first 5 minutes post-drug injection was compared across groups. In both sexes, ethanol IP resulted in increased locomotion compared to saline. Compared to control males, D2 KO males had a stronger stimulatory response to ethanol **(D)**. There was no effect of knockout on ethanol-induced locomotion in female mice **(E). (F)** Female control mice traveled greater distances following ethanol injection compared to male control mice. Data expressed as Mean ± SEM. *p<0.05; @ main effect of time; % main effect of virus; & main effect of ethanol; $ main effect of sex; # ethanol x time interaction; ^ ethanol x sex interaction; ! sex x virus interaction; ethanol x sex x virus interaction p=0.058.

### CRISPR knockout of D2Rs increases alcohol and sucrose intake in male mice

One week after the completion of ethanol-induced locomotion assay, both male (N=8/group) and female (N=6-8/group) mice were randomly assigned to either ethanol or water group. Mice in the ethanol group underwent 6 weeks of voluntary alcohol consumption in a two-bottle choice intermittent access to ethanol paradigm (Fig. 6A). Knockout of BNST D2Rs increased alcohol intake in males (Fig. 6B; [F_(1,14)_=6.33; p_virus_=0.025]; [F_(3.61,50.48)_=7.89; p_weeks_<0.001]; [F_(5,70)_=0.61; p_interaction_=0.693]) but not females (Fig. 6C; [F_(2.92,35.09)_=6.64; p_weeks_=0.001]; [F_(1,12)_=0.17; p_virus_=0.688]; [F_(5,60)_=0.45; p_interaction_=0.813]), as revealed by a two-way RM ANOVA with Geisser-Green house correction. Consistent with the literature, when compared between sexes, female mice generally consumed more alcohol than males (Fig. 6D; [F_(1,26)_=36.28; p_sex_<0.0001]) (Juárez and De Tomasi, 1999). Specifically, post-hoc analyses revealed female mice in both groups consumed more alcohol than their male counterparts ([t_26_=5.68, p_corrected_<0.0001; for controls]; [t_26_=2.95, p_corrected_=0.013; for D2 KOs]; Holm-Sidak’s correction). ([t_26_=5.68, p_corrected_<0.0001]; Holm-Sidak correction). While not significant, there was a trend toward a significant interaction between sex and virus ([F_(1,26)_=2.87; p_interaction_=0.102]; [F_(1,26)_=0.93; p_virus_=0.344]).

**Figure 6.**
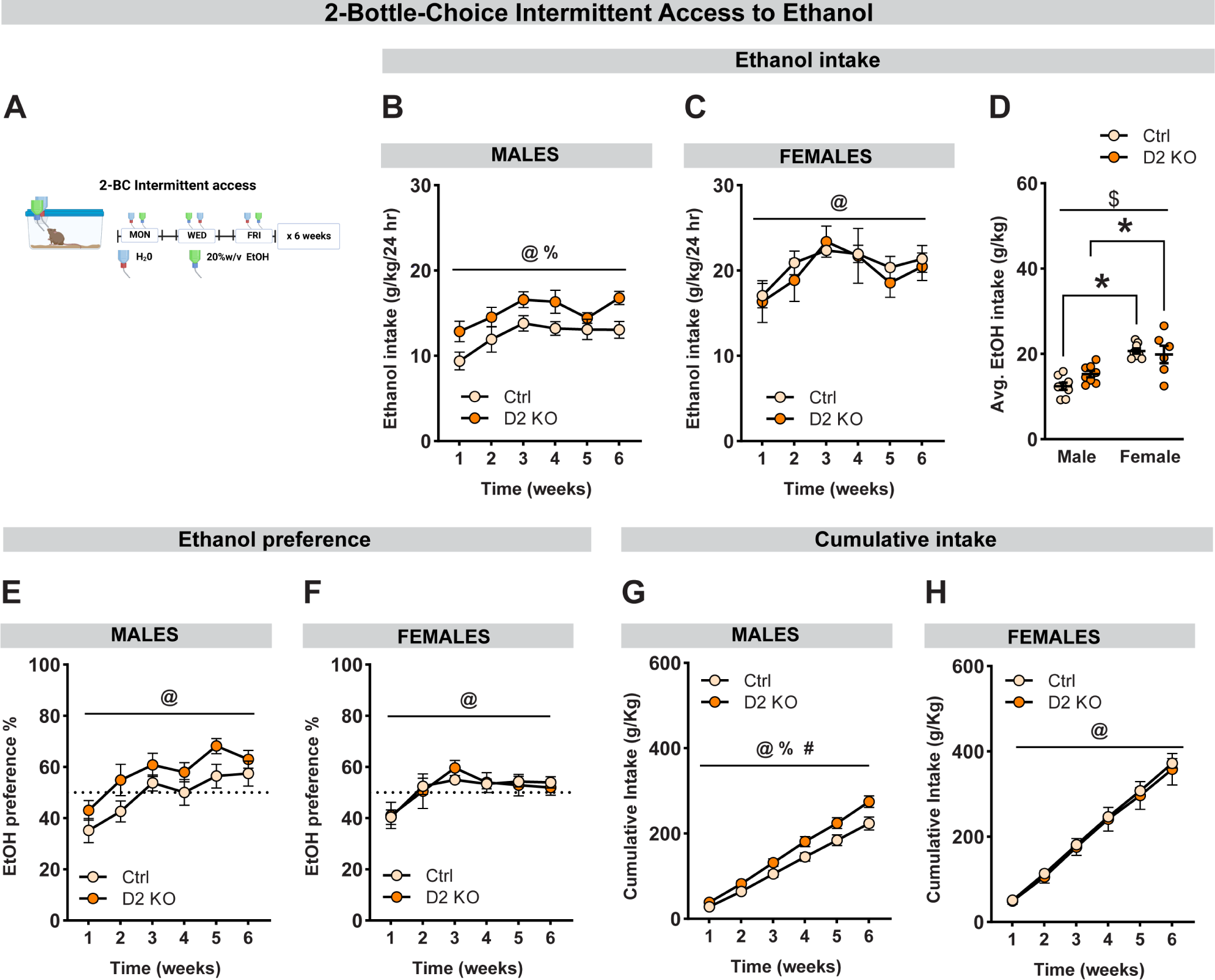
Deletion of BNST D2Rs increases alcohol consumption in males. **(A)** Schematic diagram representing two-bottle choice intermittent access to 20% w/v ethanol in mice for 6 weeks. Average weekly alcohol intake (g/kg/24h) in male **(B)** and female **(C)** mice for 6 consecutive weeks. Deletion of BNST^VGAT^ D2Rs increased alcohol intake in males but not females. **(D)** Averages of weekly intake compared between sexes. Female mice consumed more alcohol compared to male mice and there was a trend toward a sex x virus interaction (p=0.10). **(E-F)** Deletion of BNST^VGAT^ D2Rs did not affect ethanol preference in both sexes, there was a non-significant trend in increased ethanol preference in male mice (p=0.11). **(G)** Cumulative alcohol intake plotted over 6 consecutive weeks. D2 KO males had a higher cumulative intake than controls. **(H)** No effect of knockout on cumulative intake in females. Data expressed as Mean ± SEM.*p<0.05; @ main effect of time; % main effect of virus; $ main effect of sex; # virus x time interaction.

Male D2 KO mice also had a significantly higher cumulative alcohol intake (Fig. 6G; [F_(1,14)_=5.71; p_virus_=0.032]; [F_(1.25,17.47)_=576.6; p_weeks_<0.001]; [F_(5,70)_=4.91; p_interaction_=0.0007]) and a non-significant trend toward increased ethanol preference (Fig. 6E; [F_(1,14)_=2.86; p_virus_=0.113]; [F_(3.28,45.87)_=21.85; p_weeks_<0.001]; [F_(5,70)_=0.537; p_interaction_=0.748]). There was no effect of receptor deletion on average fluid intake in males (t_14_=0.74; p=0.474; data not shown). In female mice, there was a main effect of time on both cumulative intake (Fig. 6H; [F_(1.15,13.74)_=408.3; p_weeks_<0.001]) and ethanol preference (Fig. 6F; [F_(3.18,38.12)_=10.79; p_weeks_<0.001]), but no effect of either knockout ([F_(1,12)_=0.14; p_virus_=0.71; for cumulative intake]; [F_(1,12)_=0.0005; p_virus_=0.98; for ethanol preference]) or time x virus interaction ([F_(5,60)_=0.14; p_interaction_=0.983; for cumulative intake]; [F_(5,60)_=0.53; p_interaction_=0.752; for ethanol preference]. Similar to male mice, D2R knockout did not affect the mean fluid intake (t_12_=0.88; p=0.399; data not shown).

To determine the selectivity of D2R deletion effects on alcohol consumption, in one cohort, mice (N=6/group for males and N=8/group for females) were given 2 h access to a bottle of 0.5% w/v sucrose in addition to water five days after the last IA session (Fig. 7). Mice in both the ethanol and the water groups were given access to sucrose and drinking data was averaged over the last three days. Similar to effects on alcohol drinking, conditional knockout of D2Rs in the BNST significantly increased sucrose intake in males (Fig. 7A; [t_10_=3.53; p=0.006]) but not females (Fig. 7B; [t_14_=1.17; p=0.261]). A two-way ANOVA revealed a main effect of sex [F_(1,24)_=25.57; p_sex_<0.0001] but not virus [F_(1,24)_=1.48; p_virus_=0.235] on sucrose intake (Fig. 7C) along with a significant interaction between sex and deletion effects ([F_(1,24)_=8.69; p_interaction_=0.007]). Post-hoc analyses showed female controls consumed more sucrose than male controls ([t_24_=5.66; p_corrected_<0.0001 for controls]; [t_24_=1.49; p_corrected_=0.275 for KOs]) but only male D2 KO mice consumed more compared to male controls ([t_24_=2.75; p_corrected_=0.022 for males]; [t_24_=1.32; p_corrected_=0.358 for females]).

**Figure 7.**
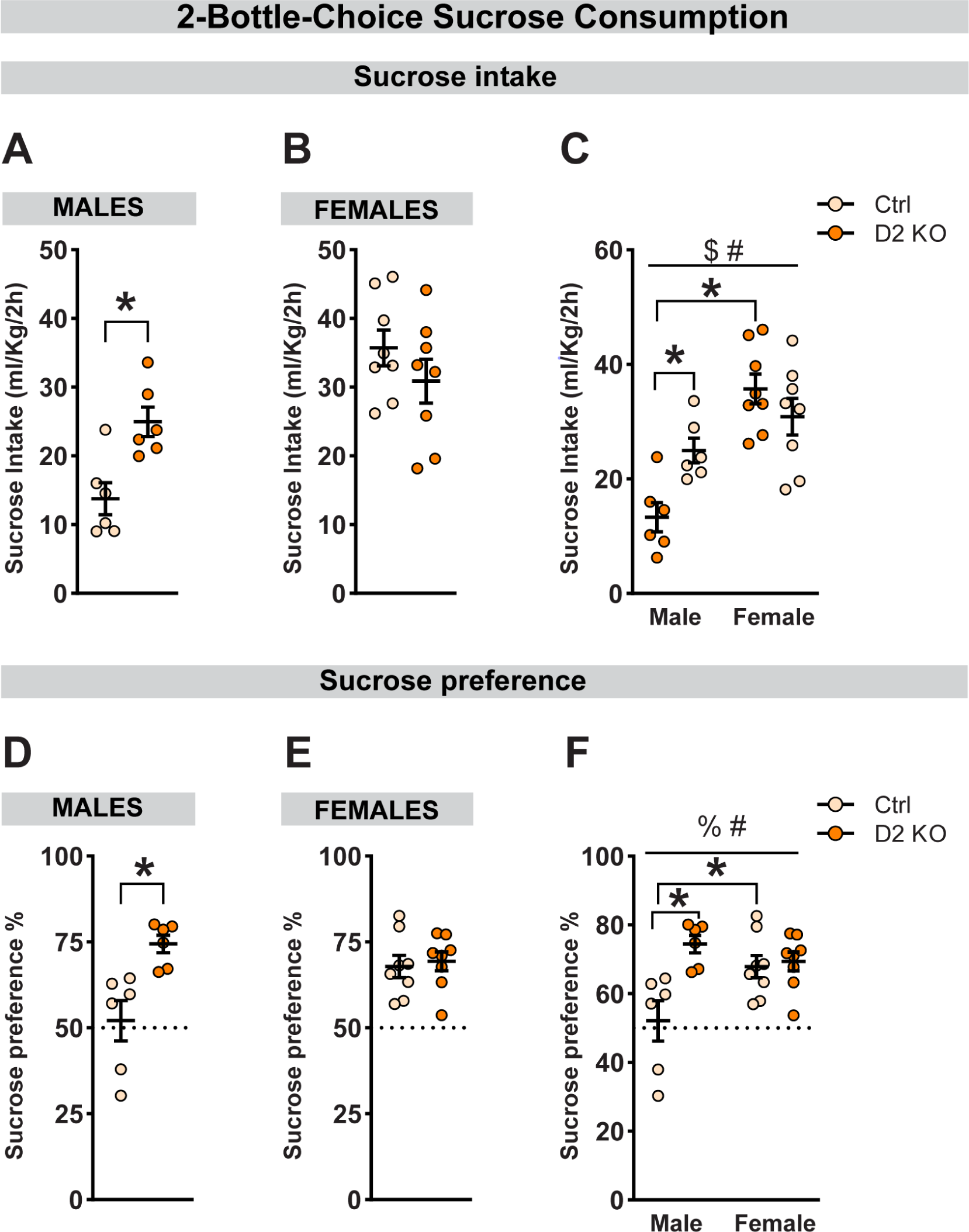
Knockout of BNST D2Rs increases sucrose intake in a sex-dependent manner. Average sucrose intake (ml/kg/2h) in male **(A)** and female **(B)** mice. Deletion of BNST^VGAT^ D2Rs increased sucrose intake selectively in male mice. **(C)** Averages of sucrose intake compared between sexes. Female controls consumed more sucrose compared to male controls. **(D-F)** Deletion of BNST^VGAT^ D2Rs also increased sucrose preference in males but not females. Female control mice had higher sucrose preference compared to male controls. There was no effect of virus in female mice. Data expressed as Mean ± SEM.*p<0.05; % main effect of virus; $ main effect of sex; # virus x sex interaction.

D2 KO male mice also had a higher sucrose preference (Fig. 7D; [t_10_=3.45; p=0.006]) which was absent in female mice (Fig. 7E; [t_14_=0.35; p=0.73]). A two-way ANOVA revealed a main effect of virus [F_(1,24)_=10.36; p_virus_=0.004] but not sex [F_(1,24)_=2.09; p_sex_=0.161] on sucrose preference (Fig. 7F) along with a significant interaction between sex and deletion effects ([F_(1,24)_=7.90; p_interaction_=0.01]). Post-hoc analyses showed female controls had higher sucrose preference compared to male controls ([t_24_=3.01; p_corrected_=0.012 for controls]; [t_24_=0.97; p_corrected_=0.57 for KOs]), but only male D2 KO mice displayed higher sucrose preference over their male counterparts ([t_24_=3.99; p_corrected_=0.001 for males]; [t_24_=0.31; p_corrected_=0.942 for females]).

## Discussion

In our current study, we used a combinatorial genetic and CRISPR/SaCas9-based approach to interrogate the contributions of BNST dopamine D2 receptors in alcohol consumption and other relevant behaviors in male and female mice. Reduction of D2R expression in BNST GABA neurons resulted in distinct sex-specific behavioral changes. Our results add to the growing body of literature supporting the interpretation that D2Rs are critical for the rewarding and locomotor properties of alcohol.

### BNST D2Rs and pain sensitivity

Preclinical studies have highlighted the role of BNST in the modulation of pain-related affect (Deyama et al., 2008; Hagiwara et al., 2013; Minami, 2019; Yu et al., 2021a). In a study by Hagiwara et al., direct injection of a D1 receptor antagonist in the BNST enhanced formalin-induced hyperalgesia only in female rats (Hagiwara et al., 2013), supporting a role for BNST DA signaling in pain. Prior work from our group identified a novel vlPAG/DR to dBNST DA circuit that modulates pain responses in a sex-specific manner (Li et al., 2016; Yu et al., 2021b). We demonstrated that activation of vlPAG/DR dopaminergic projections to the BNST resulted in reduced pain sensitivity in male mice whereas in female mice it led to increased locomotion in a salient context. We also found sex-specific differences in BNST D2R mRNA expression with lower D2R mRNA in female mice (Yu et al., 2021b). Surprisingly, in the present study, deletion of D2Rs did not alter mechanical or thermal nociception in male mice but increased mechanical nociceptive sensitivity in female mice. Our results suggest that D2R-expressing BNST GABA neurons may contribute to acute mechanical nociception.

### BNST D2Rs and ethanol-induced locomotion

In rodents, IP injection of ethanol results in a transient increase in locomotion in a dose-dependent manner (Liljequist et al., 1981; Cohen et al., 1997; Palmer et al., 2003). Enhanced sensitivity to the stimulatory effects of ethanol and resilience to the sedative properties are known to be associated with vulnerability for AUD (Erblich and Earleywine, 2003; King et al., 2011). In our present study, we observed heightened ethanol-induced stimulation in D2R KO male mice. Our findings are consistent with prior work evaluating the effects of D2R deletion on ethanol locomotion. For example, in a study by Palmer et al., mice with global deletion of D2Rs showed enhanced ethanol-induced stimulation when tested in a familiar environment (Palmer et al., 2003). Similarly, using a transgenic approach, Bocarsly et al., found mice with selective deletion of D2Rs on striatal medium spiny neurons had a similar phenotype (Bocarsly et al., 2019). Interestingly, unlike the other two studies, we found a sex-specific effect since D2R KO did not alter ethanol stimulation in female mice. Collectively, our results suggest a causal link between BNST^VGAT^ D2R downregulation and ethanol-induced stimulation. Additional work is needed to examine whether BNST D2R knockout also provides resilience to the sedative properties of acute ethanol.

### BNST D2Rs and voluntary alcohol intake

Though equivocal, it has been suggested that low levels of D2R expression is a predisposing factor for AUD (Hietala et al., 1994; Volkow et al., 1996, 2002). Thus, one might predict that reducing D2R activity either through pharmacological or genetic approaches would enhance ethanol consumption. Consistent with this idea, here we report that genetic inactivation of BNST^VGAT^ D2Rs increased voluntary alcohol drinking in male mice in a two-bottle choice intermittent access paradigm. D2R knockout also resulted in increased sucrose drinking in male mice suggesting a general role for BNST D2Rs in the general consumption of palatable reward. Interestingly, neither alcohol nor sucrose intake was affected in females, hinting toward a plausible sexually dimorphic regulation of drinking behaviors by BNST D2Rs. However, our findings differ from prior work involving pharmacological and genetic manipulations of D2Rs. For example, previous studies using D2R-deficient mice showed a reduction in ethanol preference (Phillips et al., 1998), ethanol-induced conditional place preference (Cunningham et al., 2000) as well as a lack of operant ethanol self-administration (Risinger et al., 2000). Similarly, results from studies using pharmacological methods have been inconclusive. In some studies, D2-like antagonists attenuated ethanol intake in rats and mice (Dyr et al., 1993) while in another study, D2-like antagonists increased alcohol drinking (Levy et al., 1991; Hodge et al., 1997). Specific to BNST, prior work from Eiler II et al., demonstrated pharmacological blockade of D1-like receptors but not D2-like receptors in the BNST reduced ethanol and sucrose self-administration in male and female alcohol-preferring P rats (Eiler II et al., 2003). This discrepancy with earlier work could be due to a multitude of reasons including lack of specificity, differences in animal models, and differences in experimental protocols.

### Conclusions and functional implications

Using a combination of techniques, we and others have established the role of BNST in the maintenance of AUD. For example, alcohol withdrawal drives neuroplastic changes in BNST (Francesconi et al., 2009; Kash et al., 2009; Wills et al., 2012; Pati et al., 2020) and chemogenetic manipulation of specific subpopulations of neurons alters binge-like drinking (Pleil et al., 2015; Rinker et al., 2017). Alcohol is also associated with sexually dimorphic responses within the BNST(Levine et al., 2021; Marino et al., 2021). In the current study, by combining transgenic mice with a CRISPR-based loss of function approach, we were able to dissect the contributions of D2R-expressed on BNST GABA neurons in alcohol-related behaviors. Our findings further implicate BNST in alcohol abuse by providing evidence for a causal link between BNST D2Rs and alcohol-related behaviors. Notably, though our findings suggest BNST D2Rs modulate alcohol and sucrose intake selectively in males, we cannot exclude the possibility that a lack of effect in females could be related to higher variability in female drinking data or variability in the viral expression and spread.

Previously, we and others have shown the presence of mostly non-overlapping D1 and D2 receptor-expressing neurons, primarily within the lateral part of dBNST (De Bundel et al., 2016; Melchior et al., 2021; Yu et al., 2021b). Therefore, we cannot exclude the possibility of compensatory adaptations in BNST D1R function to overcome the lower levels of D2Rs. Recent work from the Alvarez lab identified an upregulation of striatal D1R function in response to selective deletion of D2Rs from medium spiny neurons as the driving force for escalated drinking and heightened ethanol stimulation (Bocarsly et al., 2019). Future studies would be required to examine how the D1 and D2 systems work together within the BNST to modulate alcohol-relevant behaviors.

## Acknowledgments

This work was funded by the National Institutes of Health grants R01NS122230 (TLK), R21AA027460 (TLK), R01AA019454 (TLK), U01AA020911 (TLK), R01DA044315 (LSZ), and P30DA048736 (LSZ).

---

**Supplementary to Figure 1.**
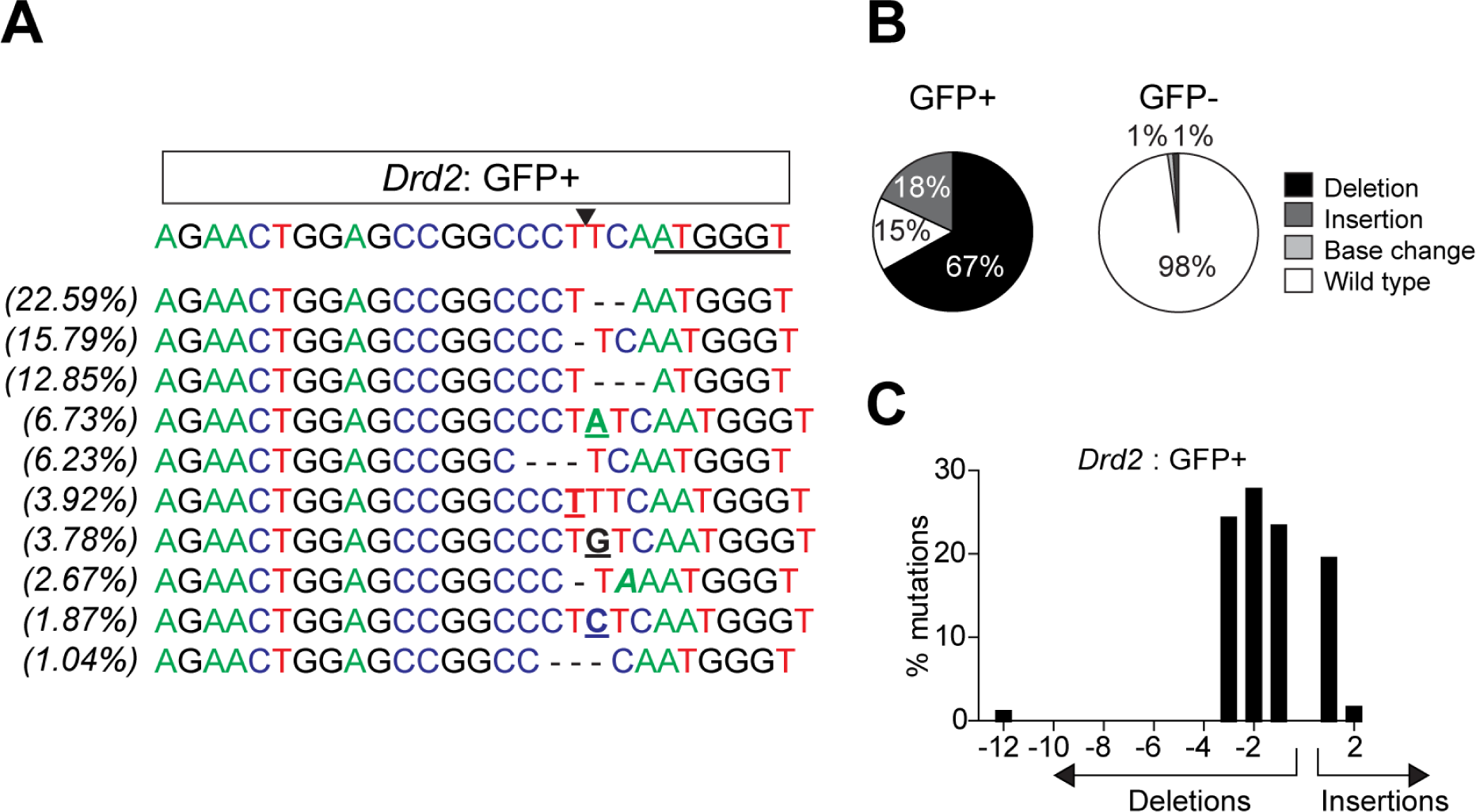
Analysis of targeted DrD2 mutagenesis in DAT-cre mice. **(A)** Analysis of FACS-sorted GFP+ nuclei from mice co-injected with AAV1-FLEX-SaCas9-sgDrd2 and AAV1-FLEX-EGFP-Kash into the VTA. sgDrd2 sequence with PAM underlined and SaCas9 cut site indicated by a black arrow. Top 10 mutations at the cut site with the percent of total reads for which they occur on the left. Base changes: bolded. Insertions: underlined. Deletions: marked with a “-” (dash). **(B)** Percentage of sequence reads with wild-type, deletions, insertions, and base changes for Drd2 in GFP+ (left) and GFP-nuclei (right). **(C)** Frequency distribution of insertions and deletions in Drd2 from GFP+ nuclei.

